## Supplementary material for "The veterinary anti-parasitic selamectin is a novel inhibitor of the mycobacterial DprE1 enzyme": Fig. S1

**SUPPORTING INFORMATION**

**The veterinary anti-parasitic selamectin is a novel enzymatic inhibitor of the *Mycobacterium tuberculosis* DprE1 protein**

Ezquerra-Aznárez, J.M. *et al.*

**Table S1. Oligonucleotides used for DprE1 mutagenesis**

| **Name** | **Sequence** |
| --- | --- |
| DprE1 C394A For | CGATCCCGGGCTGGAACGTGGCCGTGGACTTCCCGATCAAGG |
| DprE1 C394A Rev | CCTTGATCGGGAAGTCCACGGCCACGTTCCAGCCCGGGATCG |
| DprE1 C394G For | CGATCCCGGGCTGGAACGTGGGCGTGGACTTCCCGATCAAG |
| DprE1 C394G Rev | CTTGATCGGGAAGTCCACGCCCACGTTCCAGCCCGGGATCG |
| DprE1 C394S For | GATCCCGGGCTGGAACGTGTCCGTGGACTTCCCGATCAAGG |
| DprE1 C394S Rev | CCTTGATCGGGAAGTCCACGGACACGTTCCAGCCCGGGATC |
| DprE1 L282F For | CGCCGCAACTGCTCACGTTTCCGGACATCT |
| DprE1 L282F Rev | AGATGTCCGGAAACGTGAGCAGTTGCGGCG |
| DprE1 L282V For | CGCCGCAACTGCTCACGgTgCCGGACATCT |
| DprE1 L282V Rev | AGATGTCCGGcAcCGTGAGCAGTTGCGGCG |

**Table S2. Oligonucleotides used for *M. smegmatis* recombineering.** Point mutations introduced with mutagenic oligos are highlighted

| **Name** | **Sequence** |
| --- | --- |
| rpsL+ | GCGACCTTCCGGAGCGCCGAGTTCGGCTTCCTCGGAGTGGTGGTGTAAACGCGCGTGCACA |
| L282F | AGCTGCAGAAGGATCCACTGAAATTCGATGCGCCGCAACTGCTCACGTTTCCGGACATCTTCCCGAACGGCCTGGCCAACAAGTTCACGTTCATGCCGAT |
| L282V | AGCTGCAGAAGGATCCACTGAAATTCGATGCGCCGCAACTGCTCACGGTTCCGGACATCTTCCCGAACGGCCTGGCCAACAAGTTCACGTTCATGCCGA |
| dprE1-seq-F | gtgagcctggaccagttgatgaaagc |
| dprE1-seq-R | tacagcctgccaccgaactcc |

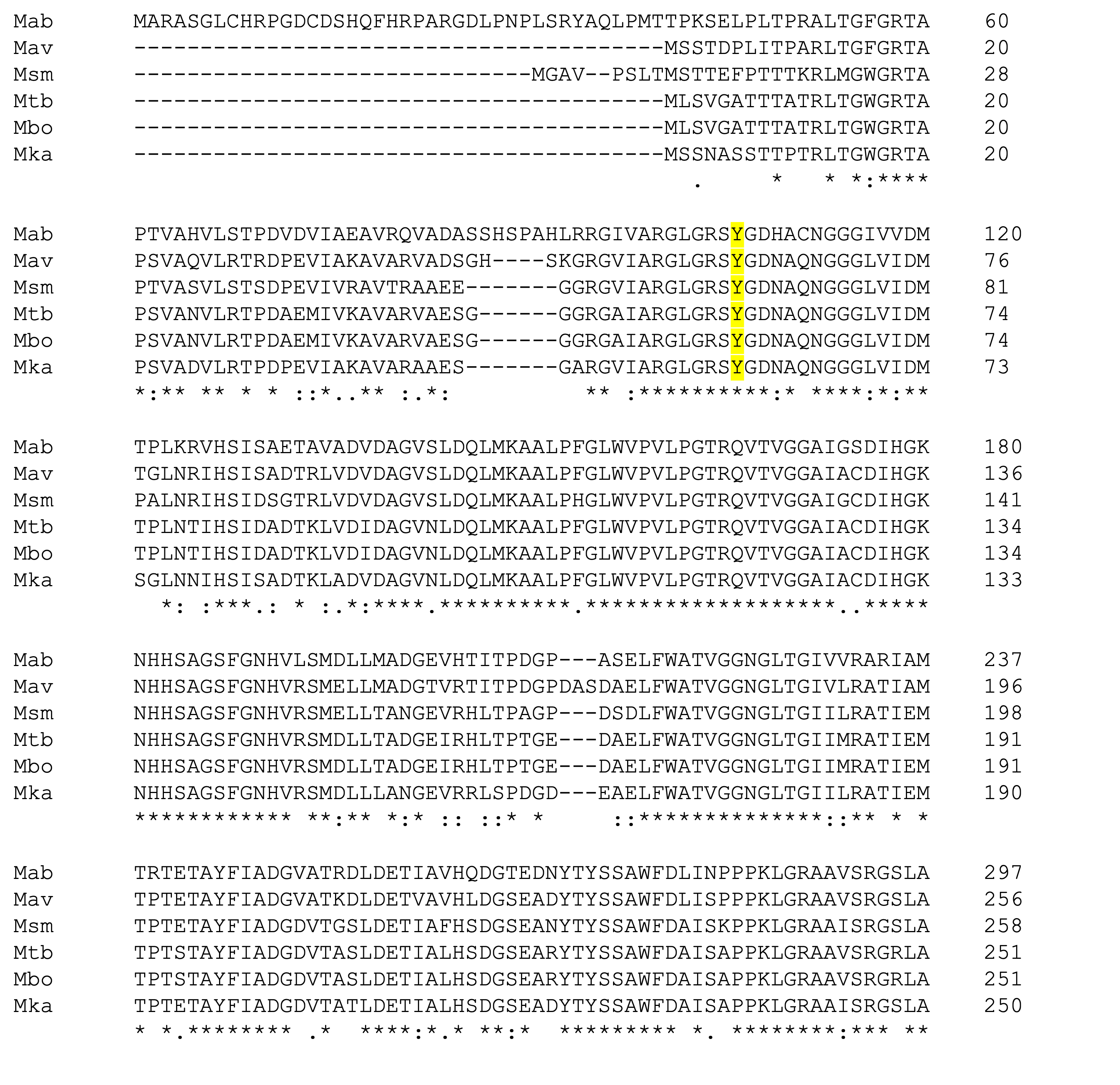

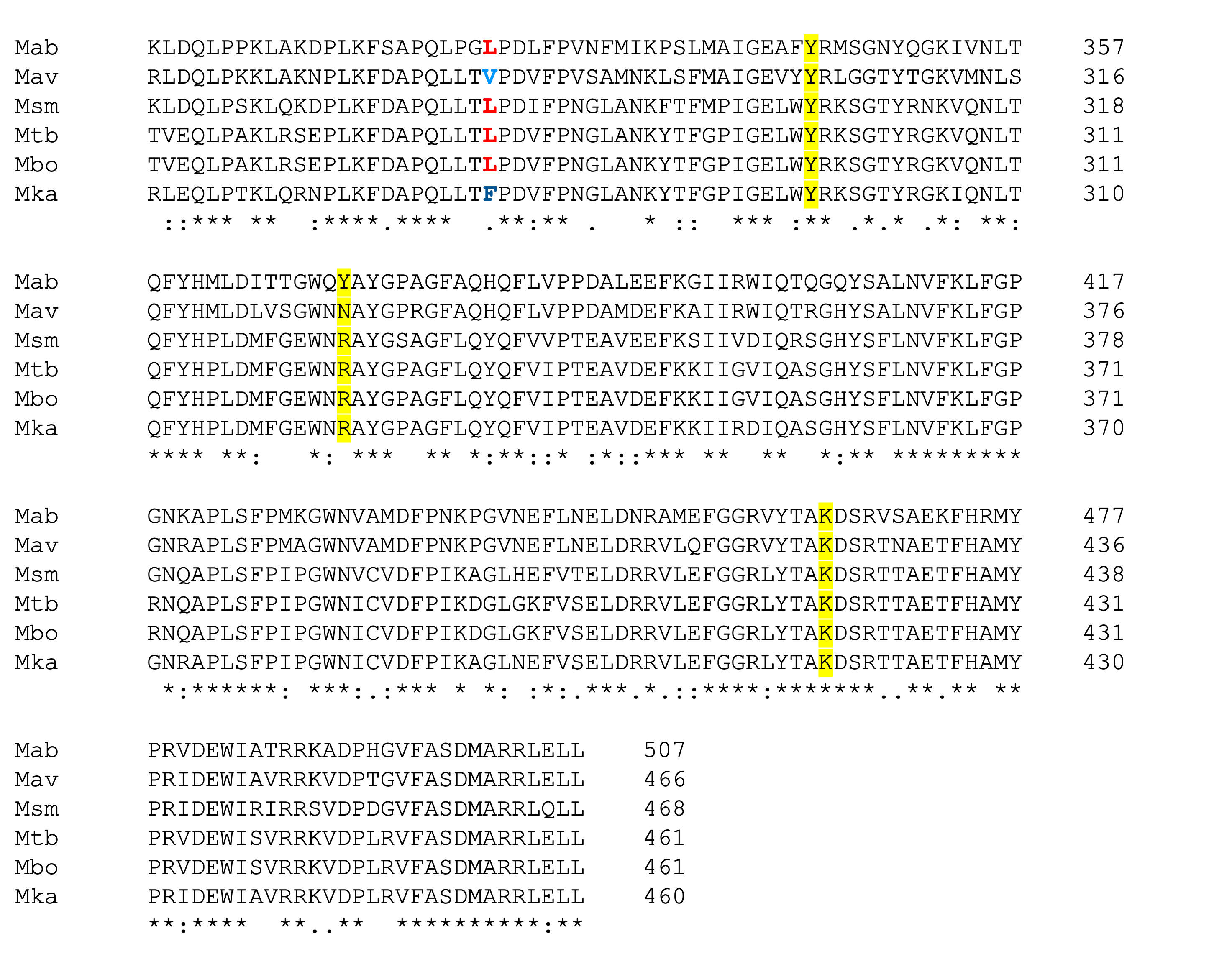

**Figure S1. Sequence alignment of mycobacterial DprE1.** Residues predicted to be relevant for selamectin binding to DprE1 are highlighted in yellow (Leu275 is shown in red) *Mab:* *Mycobacterium abscessus*; *Mav*: *Mycobacterium avium*; *Msm*: *Mycobacterium smegmatis*; *Mtb*: *Mycobacterium tuberculosis*; *Mbo*: *M. bovis*; *Mka*: *M. kansasii*

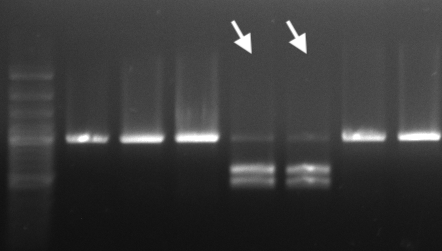

**Figure S2. Screening of *M. smegmatis* recombinants for point mutations.** Following transformation with the mixture of mutagenic oligonucleotides targeting both *dprE1* and *rpsL*, colonies isolated on streptomycin-containing were screened for the presence of point mutations in *dprE1*. A 944 bp fragment of *dprE1* was amplified by PCR, and then digested with *Kpn*2I. Wild-type *M. smegmatis* shows the undigested 959 bp product, while mixed populations (white arrows) show the undigested fragment and the two digestion products (525 and 419 bp). Left lane: Gene Ruler 100 bp Plus Ladder (ThermoFisher Scientific).

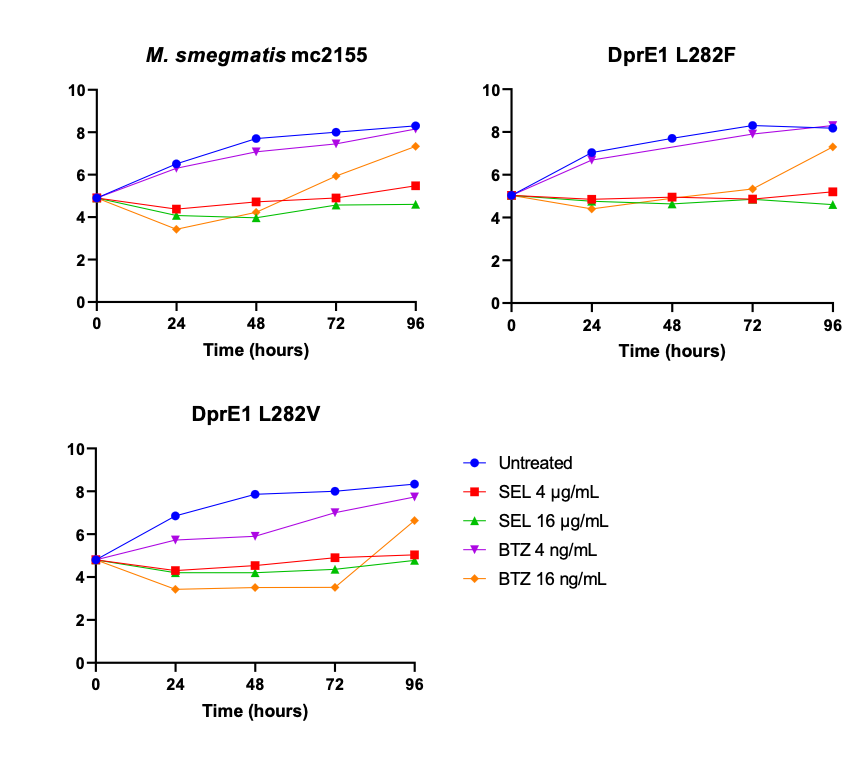

**Figure S3. Time-kill kinetics of *M. smegmatis* DprE1 point mutants.**
